# Vorinostat (SAHA) alters the skeletal muscle differentiation program

**DOI:** 10.64898/2026.05.27.727835

**Authors:** Veronica Sian, Andreas Roos, Andreas Hentschel, Jaakko Sarparanta, Per Harald Jonson, Sergio Valente, Antonello Mai, Lucia Altucci, Bjarne Udd, Angela Nebbioso, Marco Savarese

## Abstract

Epigenetic regulation, particularly histone acetylation, plays a critical role in skeletal muscle differentiation by modulating gene expression programs without altering DNA sequence. Histone deacetylases (HDACs) tightly regulate myogenesis by controlling the timing of differentiation. Pharmacological inhibition of HDACs has shown context-dependent effects on muscle cells. We investigated the effects of 1 µM SAHA (suberoylanilide hydroxamic acid) on C2C12 and L6 myoblasts during differentiation using morphological, immunofluorescence, transcriptomic, and proteomic analyses. SAHA delayed early differentiation, reducing myotube formation with partial recovery at later stages. Transcriptomic analysis revealed time-dependent changes in pathways related to cytoskeleton, cell cycle, and chromatin regulation. Proteomics showed increased mitochondrial metabolism and reduced cytoskeletal components in C2C12 cells, while L6 cells displayed alterations in muscle structural and extracellular matrix proteins. SAHA induces stage- and model-dependent reprogramming of myogenesis, highlighting the importance of timing and cellular context in HDAC-targeted therapies.

## Introduction

Epigenetic regulation is a fundamental mechanism controlling gene expression without altering the underlying DNA sequence (1). Through dynamic modifications of chromatin, such as DNA methylation, histone acetylation and methylation, and higher-order chromatin remodeling, cells establish transcriptional programs that define lineage identity and functional specialization (2). Among these mechanisms, histone acetylation plays a central role in modulating chromatin accessibility (3). Acetylation of lysine residues on histone tails reduces nucleosome compaction and promotes recruitment of transcriptional machinery, whereas histone deacetylation condenses chromatin and restricts gene expression (3). The balance between histone acetyltransferases (HATs) and histone deacetylases (HDACs) therefore serves as a critical regulatory axis in developmental and differentiation processes (4).

Myogenesis is a highly coordinated process regulated by a complex interplay of intrinsic and extrinsic mechanisms that govern the formation of muscle fibers (5). Epigenetic regulation is essential for orchestrating the transition from proliferating myogenic progenitors to terminally differentiated myofibers (6). In this context, HDACs directly interact with myogenic transcription factors, repressing premature activation of differentiation-associated genes and maintaining proliferative states (7–9). Dynamic modulation of HDAC activity is thus required for appropriate timing of cell cycle exit, activation of structural muscle genes, and myoblast fusion (10). HDAC inhibitors are increasingly being explored for the treatment of muscle disorders, with Givinostat emerging as a leading pan-HDAC inhibitor in Duchenne muscular dystrophy (DMD) (11).

Vorinostat, also known as suberoylanilide hydroxamic acid (SAHA), is a small molecule inhibitor of HDAC activity, that prevents the removal of acetyl groups from histones (12). Although widely studied in cancer for its epigenetic effects, increasing attention has been directed toward its role in skeletal muscle biology (12–14). Inhibition of class I and II HDAC promotes myogenesis, in part through activation of MyoD and myocyte enhancer factor 2 (15,16). However, the effects of HDAC inhibitors appear to be highly stage-dependent. Exposure of undifferentiated C2C12 myoblasts to HDAC inhibitors such as butyrate (17), valproic acid (18), or trichostatin A (TSA) (19,20), followed by their removal prior to induction of differentiation, promotes myogenic differentiation. In contrast, continuous exposure during the differentiation phase impairs myotube formation, despite the activation of certain muscle-related genes (21).

The aim of this study was to investigate the effects of SAHA, focusing on how SAHA-induced HDAC inhibition modulates gene reprograming and protein expression during the myogenic differentiation process. We used two different *in vitro* skeletal muscle models, C2C12 murine myoblasts and L6 rat myoblasts. C2C12 cells exhibit a robust differentiation capacity and the ability to form mature, contractile myotubes, making them a well-established model for studying late-stage myogenesis (22). In contrast, L6 cells display a more limited differentiation potential and distinct metabolic features (23). The use of these two models allows the assessment of both conserved and model-specific responses to SAHA treatment, as well as the evaluation of how HDAC inhibition impacts myogenic progression.

## Material and Methods

### Cell culture

C2C12 murine myoblasts (ATCC, CRL-1772) and L6 rat myoblasts were cultured as previously described (24). Briefly, C2C12 cells were maintained in Dulbecco’s modified Eagle medium (DMEM, Gibco), without phenol-red and pyruvate, and supplemented with 20% fetal bovine serum (SERANA S-FBS-SA-015), 1× glutamax, and 1× penicillin/streptomycin. Cells were cultured at 37°C in a humidified 5% CO₂ atmosphere and passaged at approximately 50% confluence.

For differentiation, ultra-compliant gelatin hydrogels were prepared essentially as described (22,25). C2C12 myoblasts were seeded on gelatin hydrogels. Upon reaching confluence, myoblasts were switched to differentiation medium (DMO) containing pyruvate-free DMEM with 2% heat-inactivated horse serum, 1× L-glutamine, 1× penicillin/streptomycin and 10% Opti-MEM (22). DMO was replaced daily. Collagenase type I 1 mg/mL (Gibco) was used to collect samples at day 3, 7, and 16 of differentiation.

L6 rat myoblasts (ATCC-CRL-1458) were purchased from LGC Nordic. Cells were cultured in Dulbecco’s modified Eagle medium (DMEM, Gibco), without phenol-red and pyruvate, supplemented with 10% fetal bovine serum, 1× glutamax and 1× penicillin/streptomycin, and maintained at 37°C in 5% CO2. Cells were subcultured when they reached 50% of confluence.

For differentiation, L6 myoblasts were grown on collagen coated (Collagen bovine, type I, Corning) plates in DMO, which was replaced daily. Samples were collected after 3 and 7 days of differentiation.

### Drug treatment

SAHA was dissolved in dimethyl sulfoxide. The drug was used at a final concentration of 1 µM. Cells were treated every 48 h and media was changed every 24 h.

### Cytotoxicity assay

C2C12 and L6 cell viability after SAHA treatment was assessed by the 3-(4,5-dimethylthiazol-2-yl)-2,5-diphenyl-2H-tetrazolium bromide (MTT) assay. C2C12 and L6 myoblasts were treated with SAHA 1 µM. MTT assay was performed at different time points (24 and 48 h) after SAHA treatment, as previously described (26).

### Immunofluorescence

PFA fixed C2C12 and L6 cells were permeabilized with 0.02% Triton X-100 for 10 min at RT, washed three times with 1× PBS., and blocked with BSA 5% for 1 h at RT. Samples were incubated with primary antibodies overnight at 4°C, followed by secondary antibodies incubated for 1 h at RT protected from light. Hoechst was added for 5 min at RT. C2C12 cells were mounted with PVA mounting media, as previously described (22,24). Zeiss Axio Imager M2 (Carl Zeiss AG, Oberkochen, Germany) was used for the image acquisitions.

### Total protein extraction and western blotting

Total protein extraction and western blotting were performed as previously described (24). Briefly, C2C12 and L6 cell pellets were collected and washed three times with 1× PBS and centrifuged at 1500 g for 5 minutes at 4°C. Cell pellets were resuspended in RIPA Buffer (NP-40 1%, 50 mM Tris-HCl pH 8.0, 150 mM NaCl, 10% glycerol, 1 mM EDTA) with 1x Halt™ Protease Inhibitor Cocktail (Thermo Scientific™), vortexed three times every 10 minutes, and then centrifuged at 15000 g for 20 min at 4°C. Protein concentration was determined by Bradford assay (Bio-Rad, California, US).

Protein samples were separated in TGX minigels (Bio-Rad, Hercules, CA, USA) and transferred on nitrocellulose membranes with the Trans-Blot Turbo system (Bio-Rad). Total protein was stained with Revert 520 Total Protein Stain (Li-Cor Biosciences, Lincoln, NE, USA) and scanned with Odyssey M scanner (Li-Cor). The stain was detected in the 520 nm channel for quantification and normalization analyses.

Membranes were blocked with 5% milk in PBS for 1 h, and washed with 1× PBS + 0.1% Tween (PBS-T). Primary antibodies were diluted in 1% milk and incubated at 4°C overnight, followed by fluorescent secondary antibodies (1:10000) diluted in milk 5% for 1 hour at RT. Primary antibodies used are the following: anti-Nat10 (Abcam ab251186), anti-MyHC (DSHB), anti-Mybpc1 (Abcam ab55559), anti-Calsequestrin 1/2 (Abcam ab3516), anti-Troponin T (Novocastra). Odyssey scanner was used to detect the signals. Empiria studio (Li-Cor) was used for the analysis. All quantitative data are expressed as mean ± standard error of the mean.

### Histone extraction and western blotting

Histone extraction was performed as reported earlier (24). C2C12 samples were collected 3 days after the switch to DMO, to capture early myogenic dynamics, and after 7 days and 16 days, representing an intermediate and late stage of differentiation, respectively. L6 samples were collected 7 days after the switch to DMO. Cell pellets were washed with 1× PBS, then resuspended in Triton Extraction Buffer (TEB: PBS containing 0.5% Triton X-100 (v/v), 2 mM phenylmethylsulfonyl fluoride (PMSF), 0.02% (w/v) NaN3) and kept on the rotator for 10 minutes. Then, samples were centrifuged at 2000 g for 10 minutes at +4°C. Supernatant was removed and the pellet washed in half the volume of TEB and centrifuged as before. Pellet was resuspended in 0.2N HCl for acid extraction over night at 4°C, followed by centrifugation at 2000 rpm for 10 minutes at 4°C. Protein content was determined using Bradford assay. Western blot analyses were performed for three acetylation targets, namely histone 3 lysine 9/14 acetylated (H3K9/14ac, Cell signaling), histone 3 lysine 18 acetylated (H3K18ac, Diagenode), histone 4 lysine 8 acetylated (H4K8ac, Abcam). Statistical analysis was performed as previously described.

### Transcriptomic analysis

A total of 30 C2C12 samples and 20 L6 samples were processed for transcriptome sequencing (Supplementary Table 1-2). Total RNA was extracted with Trizol (Invitrogen). Transcriptomic was performed as mentioned previously (24). RNA concentration and purity was measured using NanoDrop 2000 (Thermo Fisher Scientific, Wilmington, DE). RNA integrity was assessed using the RNA Nano 6000 Assay Kit of the Agilent Bioanalyzer 2100 system (Agilent Technologies, CA, USA). mRNA was purified from total RNA using oligo-dT-attached magnetic beads. Sequencing libraries were generated using NEBNext UltraTM RNA Library Prep Kit for Illumina (New England Biolabs, USA) following manufacturer’s recommendations.Library was sequenced using an Illumina NovaSeq6000. Library preparation and run have been performed at Biomarker Technologies (BMK) GmbH. About 20 million reads per sample were generated.

Hisat2 tools software was used to map with reference genome (Mus_musculus GRCm39 for C2C12, and GRCr8_GCF_036323735.1.genome.fa for L6 cells). Differential expression analysis was performed using DESeq2 (27). P-values were adjusted using the Benjamini and Hochberg’s approach for controlling the false discovery rate. Genes with an adjusted p-value < 0.01 were assigned as differentially expressed. BMKCloud (www.biocloud.net) and SninyGO 0.85.1 tool was used to perform Gene Ontology (GO) analysis (28). The analysis was performed with the following settings: an FDR cutoff of q = 0.01, a pathway size range of 2 to 5000, and the options “remove redundancy” and “abbreviate pathways” enabled. GO analysis of differentially expressed genes (DEGs) was performed in three categories: biological process (BP), cellular component (CC), and molecular function (MF). Sample M3, M15, and M18 were excluded from the analysis because they failed data quality control.

### Proteomics Mass Spectrometry and Data Analysis

Proteomics Mass Spectrometry was performed as previously mentioned (24). A total of 4 control samples and 4 SAHA-treated samples at day 7 of differentiation were processed for proteomics for both C2C12 and L6 cells. For data analysis, proteins identified with at least 2 unique peptides were retained for downstream analyses. Differential protein abundance between groups was assessed by calculating pairwise protein ratios followed by a t-test (p-values < 0.05). Proteins showing a fold change > 2 were classified as upregulated, whereas proteins with a fold change < 0.5 were considered downregulated. Functional enrichment of the significantly dysregulated proteins was carried out using the PANTHER (Protein ANalysis THrough Evolutionary Relationships) database. Overrepresentation analysis was applied to identify enriched Gene Ontology (GO) categories related to biological processes, molecular functions, and cellular components.

## Results

### Cell viability and morphology after SAHA treatment

We performed MTT assay to evaluate the effects of 1 µM SAHA exposure on C2C12 and L6 cell viability at 24, 48, and 72 h, respectively. SAHA was administered to the cells every 48 h. After 24 h of treatment, the drug induced a modest reduction of viability to approximately 88% in C2C12 cells and 95% in L6. At later time points, the effect became more pronounced in C2C12 cells, with viability decreasing to ∼70% at 48–72 hours. In contrast, L6 cells showed a milder response, maintaining viability at ∼83% and ∼86% at 48 and 72 hours, respectively, indicating a differential sensitivity between the two cell lines (Supplementary Figure 1).

To investigate the impact of SAHA on myogenic differentiation, C2C12 cells were differentiated on hydrogel substrates for 16 days, while L6 cells were cultured on collagen and differentiated for 7 days, reflecting their lower differentiation capacity. SAHA treatment (1 µM) was initiated 48 hours prior to the induction of differentiation and maintained throughout the experiment with administration every 48 hours, while the differentiation medium was refreshed daily. Phase-contrast imaging was performed at early (day 3), intermediate (day 7), and late (day 16) stages to monitor morphological changes during differentiation (Figure 1a–b). In C2C12 cells, SAHA treatment delayed early differentiation, as evidenced by reduced myotube formation at day 3. However, by day 7 and day 16, multinucleated myotubes were present in both control and treated conditions (Figure 1a). Notably, treated cultures exhibited a lower density of myotubes compared to controls, although spontaneous contractile activity was still observed (Supplementary video 1–2). A similar trend was observed in L6 cells, where SAHA treatment reduced myotube formation at early stages (day 3) and moderately affected differentiation at day 7 (Figure 1b). Consistent with previous observations, L6 cells displayed limited maintenance of the differentiated phenotype beyond day 7 (23).

**Figure 1.**
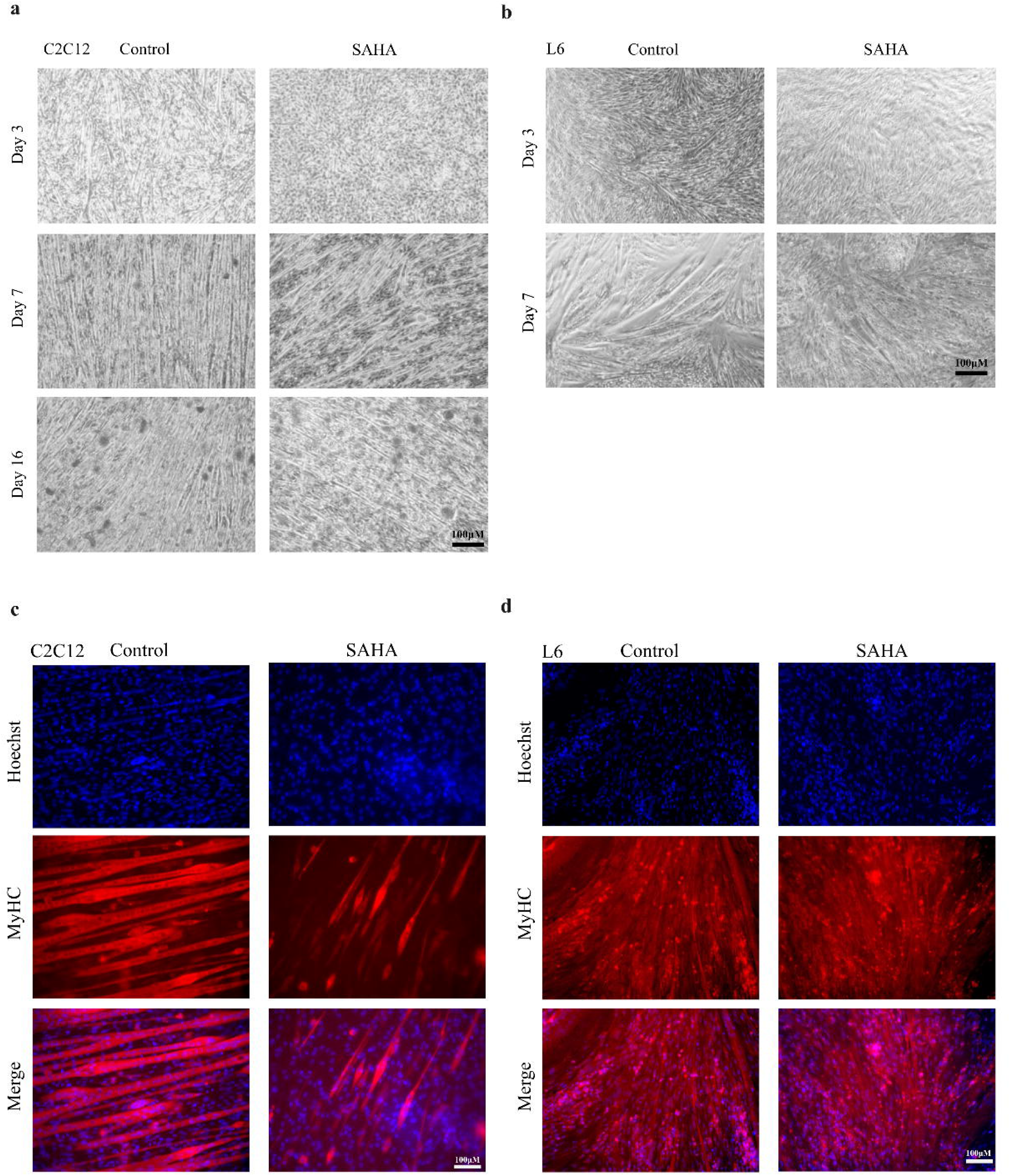
Morphological effects of SAHA on myotube maturation. (a) C2C12 and (b) L6 cells representative phase-contrast images of control and SAHA-treated cells at different stages of differentiation. For C2C12 cells images at day 3, 7, and 16 of differentiation were acquired, and for L6 cells images at day 3 and 7. Scale bar 100 μM. Immunofluorescence analyses of (c) C2C12 cells and (d) L6 cells at day 7 of differentiation. Myosin heavy chain (MyHC) in red and nuclei in blue. Images were acquired at 20× magnification. Scale bar 100 μm.

Immunofluorescence analyses for Myosin heavy chain (MyHC) at day 7 in both C2C12 and L6 cells showed reduced myotube formation and maturation compared to controls (Figure 1c-d), further illustrating the effects of SAHA on myogenic differentiation.

### Protein expression

Western blot analysis was performed to evaluate the expression of key myogenic markers and histone acetylation levels in C2C12 and L6 cells at day 3 and day 7 of differentiation following SAHA treatment (Figure 2a-b). In C2C12 cells, protein expression of Myosin heavy chain (MyHC), Myosin binding protein C1 (Mybpc1), and Troponin T1 (Tnnt1) was upregulated in the controls confirming myogenic differentiation and myotubes maturation. SAHA treatments induced a reduction in MyHC levels, while Mybpc1 and Tnnt1 levels were comparable to control conditions, as confirmed by the quantification (Figure 2a). In L6, control samples displayed a strong increase in myogenic markers at day 7, especially Mybpc1 and Tnnt1. However, SAHA treatment reduced their expression, resulting in lower protein levels compared to controls, consistent with a delay or impairment in differentiation (Figure 2b).

**Figure 2.**
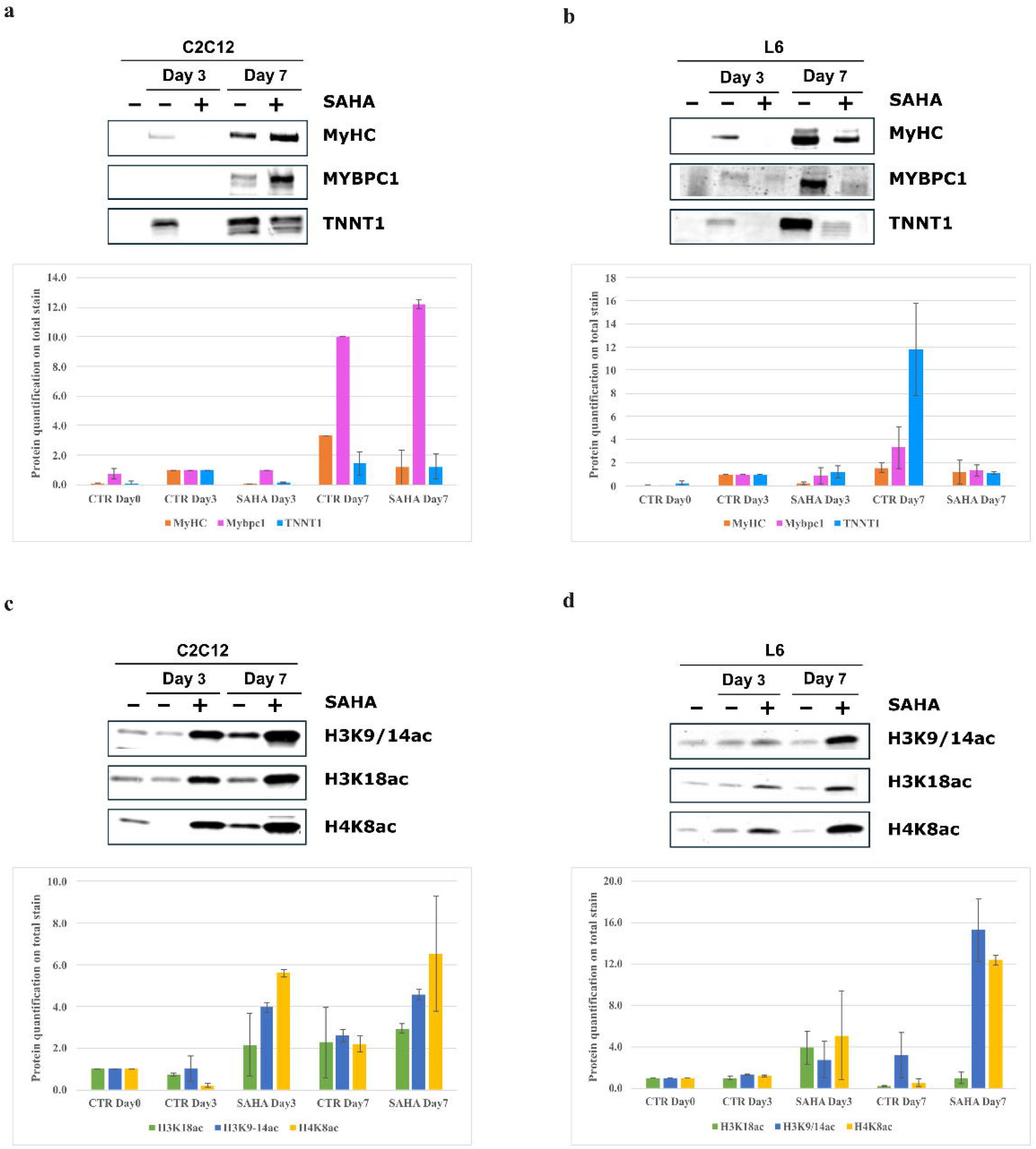
Western blot analyses of markers of differentiation and histone acetylation markers in C2C12 and L6 differentiation. a) Representative western blots of differentiation markers at day 7 of differentiation — Myosin heavy chain II (MyHC), Mybpc1, and Troponin T (Tnnt1) in control and SAHA-treated C2C12 and b) L6 cells (top). Bar chart representing the normalization of proteins on total protein stain. Control day 3 was considered as reference control (bottom). Data are representative of two independent experiments. c) Representative western blots for histone acetylation marks (H3K9/14ac, H3K18ac, H4K8ac) in C2C12 and d) L6 cells at day 0, 3, and 7 of differentiation in control and SAHA-treated cells (top). Histograms representing histone quantification normalized on total protein stain. Histone levels are expressed relative to control cells at day 0 (bottom). Data are presented as mean ± SEM (n = 2).

To assess the epigenetic effects of SAHA, histone acetylation levels were analyzed (Figure 2c-d). Western blot analyses for acetylated histones H3K18 (H3K18ac), H3K9/14 (H3K9/14ac), and H4K8 (H4K8ac) in C2C12 and L6 cells at day 3 and day 7 of differentiation confirmed the HDAC inhibitory activity of SAHA. In C2C12 cells, the level of the three histones in the control was stable at the first stage of differentiation compared to day 0 but then increased at day 7, while SAHA treatment increased acetylation levels already at day 3 of differentiation (Figure 2c). In L6 cells, acetylation level remains stable in the controls at day 3 and then diminished at day 7, while SAHA treatment increased the acetylation level of histone H3K9/14ac and H4K8ac at 3 and 7 days of differentiation compared to the control. Histone H3K18ac increased only at day 3 in the treatment group.

### Transcriptomic profile of SAHA treatment in C2C12 and L6 cells during differentiation

Transcriptomic analysis of both C2C12 and L6 cells was performed after 3 and 7 days of differentiation, and also after 16 days of differentiation on C2C12. In C2C12 cells, principal component analysis (PCA) showed a clear separation of control and SAHA-treated cells along the PC2 (Figure 3a). The PCA also showed a clear separation of the control cells at different days of differentiation underlying the transcriptomic changes that occur between myoblasts and mature myotubes (day 16), as we have also showed previously (22). Moreover, the analysis showed a clustering of treated cells at both day 7 and day 16 of differentiation.

**Figure 3.**
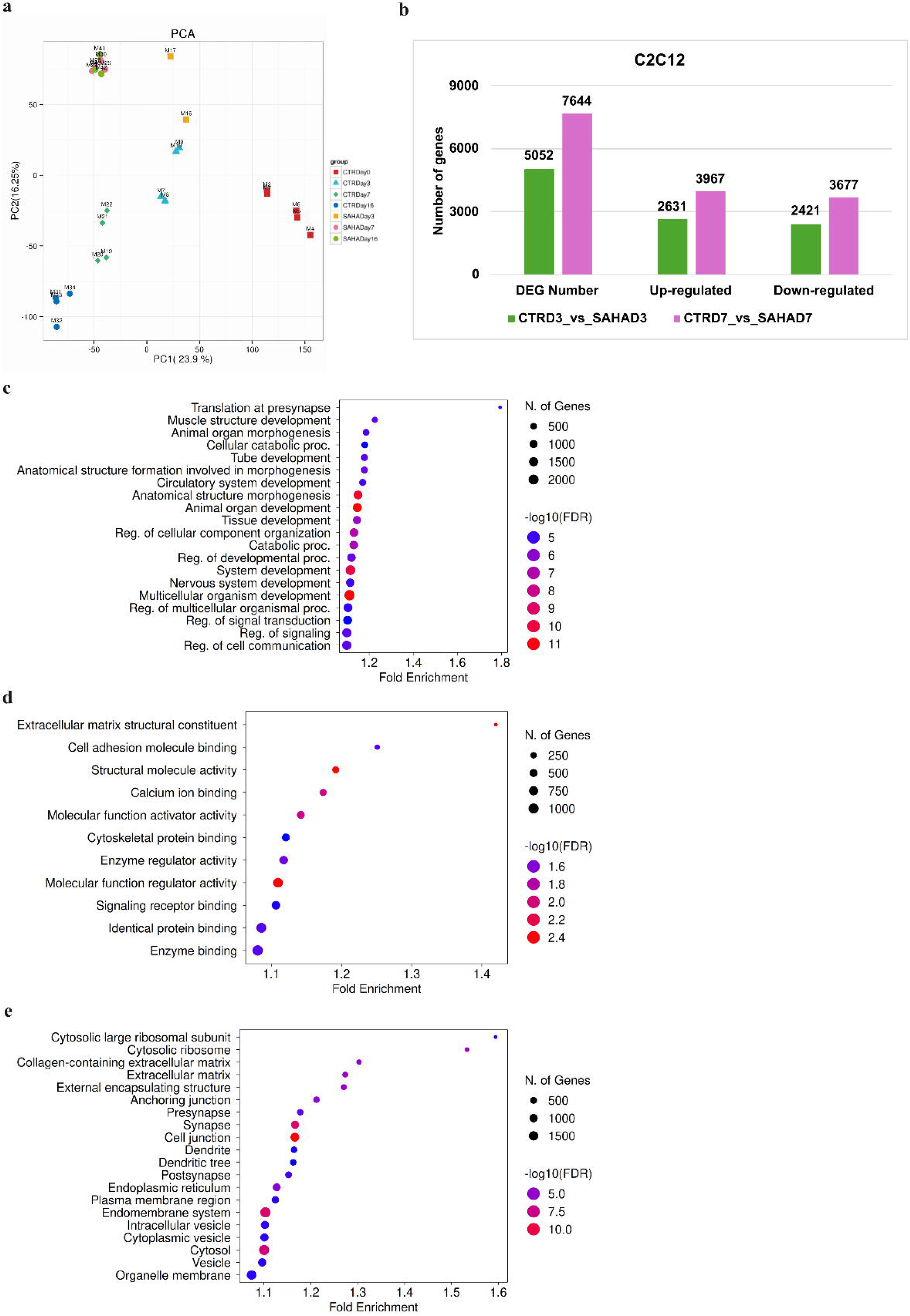
Transcriptomic analysis of C2C12 cells upon SAHA treatment. a) Two-dimensional principal component analysis (PCA) of control and SAHA-treated C2C12 samples projected along PC1 (23,9%) and PC2 (16,25%), illustrating the progressive separation of samples according to treatment and differentiation stage. b) Bar plot summarizing the number of differentially expressed genes (DEGs) at day 3 and day 7 of differentiation. Bars indicate the total number of DEGs, the up-and down-regulated in SAHA groups compared to the control. Green bars indicate the number of DEGs at day 3 of differentiation, while purple bars represent the DEGs at day 7 of differentiation. The y-axis denotes DEG counts. c) Biological process, d) Molecular function, and e) Cellular component GO databases were used to analyse the enrichment pathways in C2C12 cells at day 7 of differentiation by using ShinyGO 0.85.1 tool. FDR cutoff of q=0.01 was used for the analysis. The size of the circle represents the number of genes, with larger circles representing more genes. The colour represents the strength of the FDR, with red and blue indicating higher and lower FDR values, respectively.

Pairwise comparison between control and SAHA-treated C2C12 cells revealed a progressive increase in transcriptional changes over time. At day 3 a total of 5,052 genes were differentially expressed (2,631 were up-regulated and 2,421 were down-regulated), at day 7 a total of 7,644 (3,967 up-regulated and 3,677 down-regulated genes) (Figure 3b).

Gene ontology enrichment analysis was performed using the ShinyGO platform, based on differentially expressed genes (DEGs) identified during differentiation upon SAHA treatment. Biological process (BP), Molecular function (MF), and Cellular component (CC) database were used to analyse the most enriched pathways. At day 3, cell development, cell differentiation, and developmental processes were the most significantly enriched BP pathways (Supplementary Figure S2a). MF analysis revealed pathways predominantly associated with actin binding (Supplementary Figure 2b), while CC analysis showed muscle structural components, including sarcomere, myofibril, and contractile fiber organization (Supplementary Figure 2c).

By day 7, the enrichment profile shifted toward BP (Figure 3c) related to tissue development, morphogenesis, and regulation of cellular organization, while extracellular matrix-related terms and cell adhesion processes remained significantly MF enriched pathways (Figure 3d). CC enrichment of ribosomal and translational components suggested increased protein synthesis activity, supporting the maintenance and functional maturation of differentiated muscle cells (Figure 3e).

In L6 cells, PCA analysis performed at day 3 and day 7 of differentiation showed a clear separation between the control and the treated cells along PC2 (Figure 4a). Pairwise comparison showed that at day 3 of differentiation 6,893 genes were differentially expressed (3,509 up-regulated and 3,385 down-regulated), and at day 7 a total of 8,970 genes (4,483 up-regulated and 4,488 down-regulated) (Figure 4b).

**Figure 4.**
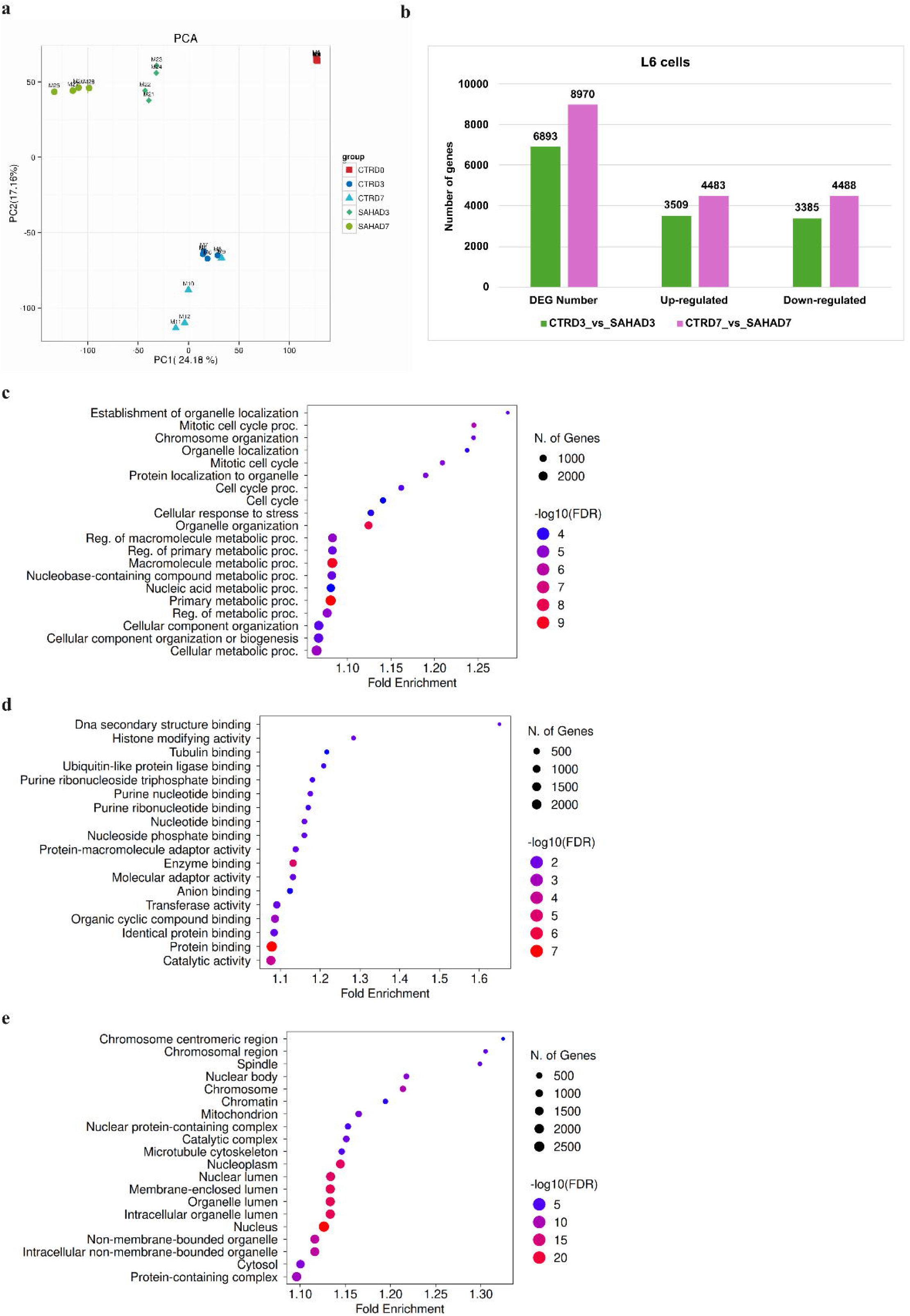
Transcriptomic profile of L6 cells upon SAHA treatment. a) Two-dimensional principal component analysis (PCA) of control and SAHA-treated L6 samples projected along PC1 (24,18%) and PC2 (17,16%), illustrating the progressive separation of samples according to treatment and differentiation stage. b) Bar plot summarizing the number of differentially expressed genes (DEGs) at day 3 and day 7 of differentiation. Bars indicate the total number of DEGs, the up- and down-regulated in SAHA groups compared to the control. Green bars indicate the number of DEGs at day 3 of differentiation, while purple bars represent the DEGs at day 7 of differentiation. The y-axis denotes DEG counts. c) Biological process, d) Molecular function, and e) Cellular component GO databases were used to analyse the enrichment pathways in L6 cells at day 7 of differentiation by using ShinyGO 0.85.1 tool. FDR cutoff of q=0.01 was used for the analysis. The size of the circle represents the number of genes, with larger circles representing more genes. The colour represents the strength of the FDR, with red and blue indicating higher and lower FDR values, respectively.

GO enrichment analysis was performed at day 3 (Supplementary Figure 3a-c) and day 7 of differentiation (Figure 3c-e). At day 3, a prominent dysregulation of pathways associated with cellular organization and proliferation was observed. In particular, BP analysis highlighted significant enrichment in terms related to mitotic progression and cell cycle regulation (Supplementary Figure 3a). MF analysis revealed enrichment in protein binding and catalytic activity (Supplementary Figure 3b). CC analysis indicated enrichment in structures associated with cell division and cytoskeletal organization (Supplementary Figure 3c).

At day 7 of differentiation, BP analysis revealed enrichment in pathways related to cell cycle, cellular component organization, and metabolic processes (Figure 4c). MF enriched pathways were mostly associated with histone-related function, tubulin binding, and catalytic activity (Figure 4d), suggesting epigenetic modulation together with cytoskeletal regulation. CC analysis revealed enrichment in chromatin, microtubule cytoskeleton, nuclear lumen, and cytosol (Figure 4e).

### Proteomic profile

To evaluate the effects of SAHA on protein abundances during myogenic differentiation in an unbiased fashion, we moreover performed proteomic analyses in both C2C12 and L6 cells at day 7 of differentiation. In C2C12 cells, a total of 195 proteins were upregulated and 272 were downregulated following SAHA treatment. To characterize the functional impact of these protein changes, GO enrichment analysis was performed using PANTHER, and the top 10 enriched terms for BP, MF, and CC were examined for both upregulated (Table 1) and downregulated protein sets (Table 2), respectively. This revealed a coordinated remodeling of metabolic pathways and cytoskeletal architecture during differentiation and moreover unveiled that SAHA treatment induced a marked upregulation of proteins associated with mitochondrial function and oxidative metabolism. This was further supported by MF pathways related to electron transfer and oxidoreductase activity, as well as CC terms related to the mitochondrial membrane and inner membrane complexes (Table 1).

**Table 1:**
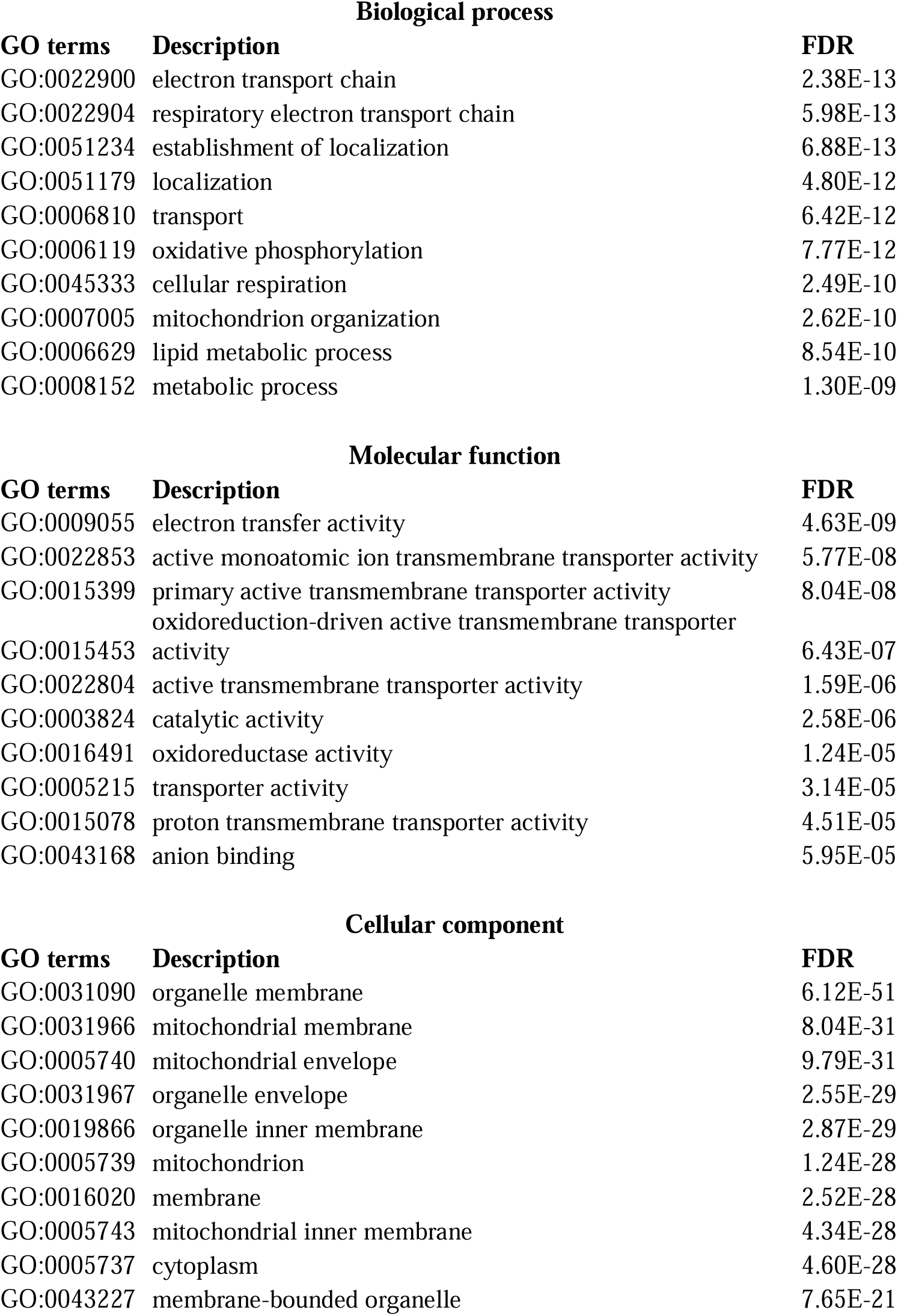
Proteomics profile of C2C12 up-regulated terms. Panther Gene Ontology (GO) analyses of proteomics data based on biological processes, molecular functions, and cellular component. The top 10 terms per each are shown.

**Table 2:** Proteomics analysis of C2C12 down-regulated terms. Panther Gene Ontology (GO) analyses of proteomics data based on biological processes, molecular functions, and cellular component. The top 10 terms per each are shown.

Biological process
| GO terms | Description | FDR |
| --- | --- | --- |
| GO:0030029 | actin filament-based process | 9.19E-14 |
| GO:0007010 | cytoskeleton organization | 5.91E-12 |
| GO:0030036 | actin cytoskeleton organization | 8.95E-12 |
| GO:0097435 | supramolecular fiber organization | 2.59E-11 |
| GO:0006996 | organelle organization | 2.40E-10 |
| GO:0016043 | cellular component organization | 4.64E-09 |
| GO:0009987 | cellular process | 4.66E-09 |
| GO:0071840 | cellular component organization or biogenesis | 9.47E-09 |
| GO:0008152 | metabolic process | 2.11E-08 |
| GO:0006006 | glucose metabolic process | 3.08E-08 |

Molecular function
| GO terms | Description | FDR |
| --- | --- | --- |
| GO:0008092 | cytoskeletal protein binding | 1.50E-18 |
| GO:0005515 | protein binding | 1.96E-18 |
| GO:0003779 | actin binding | 4.79E-18 |
| GO:0005488 | binding | 2.57E-16 |
| GO:0051015 | actin filament binding | 6.03E-10 |
| GO:0038023 | signaling receptor activity | 6.04E-10 |
| GO:0060089 | molecular transducer activity | 7.25E-10 |
| GO:0004888 | transmembrane signaling receptor activity | 1.12E-09 |
| GO:0004930 | G protein-coupled receptor activity | 1.02E-08 |
| GO:0044877 | protein-containing complex binding | 4.58E-08 |

Cellular component
| GO terms | Description | FDR |
| --- | --- | --- |
| GO:0005737 | cytoplasm | 9.82E-36 |
| GO:0005829 | cytosol | 1.17E-31 |
| GO:0005622 | intracellular anatomical structure | 2.11E-27 |
| GO:0043292 | contractile muscle fiber | 6.38E-24 |
| GO:0030016 | myofibril | 2.18E-23 |
| GO:0030017 | sarcomere | 2.88E-20 |
| GO:0099081 | supramolecular polymer | 2.74E-19 |
| GO:0099512 | supramolecular fiber | 3.13E-19 |
| GO:0099080 | supramolecular complex | 1.91E-17 |
| GO:0043229 | intracellular organelle | 7.52E-16 |

Downregulated proteins were highly enriched in actin filament-based processes, cytoskeleton organization, and supramolecular fiber assembly, supported by molecular functions such as actin binding and cytoskeletal protein binding, and cellular components including myofibrils and sarcomeres. This pattern is consistent with the reduced myotube maturation observed morphologically and suggests an extensive reorganization of structural elements (Table 2).

Unbiased proteomic profiling of L6 at day 7 of differentiation showed 49 up-regulated and 82 down-regulated proteins after SAHA treatment. GO enrichment analysis was performed using PANTHER to unravel affected BP, MF and CC. This revealed a strong impact of increased proteins on pathways associated with muscle function and maturation among upregulated proteins. BP such as actin–myosin filament sliding, and muscle structure development were highly significant, supported by MF including actin binding and cytoskeletal motor activity, and CC such as myofibrils, sarcomeres, and myosin complexes. This indicates an advanced structural maturation and functional specialization of L6 myotubes (Table 3).

**Table 3:** Proteomics profile of L6 up-regulated terms. Panther Gene Ontology (GO) analyses of proteomics data based on biological processes, molecular functions, and cellular component. The top 10 terms per each are shown.

Biological process
| Go terms | Description | FDR |
| --- | --- | --- |
| GO:0006936 | muscle contraction | 2.46E-07 |
| GO:0003012 | muscle system process | 2.97E-06 |
| GO:0032501 | multicellular organismal process | 2.32E-04 |
| GO:0009888 | tissue development | 4.84E-04 |
| GO:0033275 | actin-myosin filament sliding | 5.06E-04 |
| GO:0048731 | system development | 5.41E-04 |
| GO:0061061 | muscle structure development | 5.73E-04 |
| GO:0048856 | anatomical structure development | 6.11E-04 |
| GO:0006941 | striated muscle contraction | 6.81E-04 |
| GO:0030049 | muscle filament sliding | 7.24E-04 |

Molecular function
| Go terms | Description | FDR |
| --- | --- | --- |
| GO:0003779 | actin binding | 3.69E-08 |
| GO:0005515 | protein binding | 9.80E-05 |
| GO:0005488 | binding | 1.31E-04 |
| GO:0008092 | cytoskeletal protein binding | 1.99E-04 |
| GO:0051015 | actin filament binding | 3.49E-04 |
| GO:0000146 | microfilament motor activity | 1.07E-03 |
| GO:0003774 | cytoskeletal motor activity | 3.03E-03 |
| GO:0044877 | protein-containing complex binding | 9.99E-03 |

Cellular component
| Go terms | Description | FDR |
| --- | --- | --- |
| GO:0043292 | contractile muscle fiber | 1.02E-14 |
| GO:0030016 | myofibril | 4.46E-12 |
| GO:0015629 | actin cytoskeleton | 2.28E-09 |
| GO:0016459 | myosin complex | 1.27E-08 |
| GO:0099081 | supramolecular polymer | 4.51E-08 |
| GO:0099512 | supramolecular fiber | 5.42E-08 |
| GO:0032982 | myosin filament | 2.99E-07 |
| GO:0030017 | sarcomere | 5.73E-06 |
| GO:0005856 | cytoskeleton | 6.50E-06 |
| GO:0016460 | myosin II complex | 2.37E-05 |

In contrast, downregulated proteins were primarily associated with extracellular matrix (ECM) organization and collagen-related processes, including collagen fibril organization and extracellular structure development (Table 4). This suggests that SAHA reduces ECM-related and connective tissue-like features in L6 cultures.

**Table 4:** Proteomics analysis of L6 down-regulated terms. Panther Gene Ontology (GO) analyses of proteomics data based on biological processes, molecular functions, and cellular component. The top 10 terms per each are shown.

Biological process
| Go terms | Description | FDR |
| --- | --- | --- |
| GO:0006564 | L-serine biosynthetic process | 8.10E-04 |
| GO:0009070 | serine family amino acid biosynthetic process | 1.00E-03 |
| GO:0030199 | collagen fibril organization | 1.30E-03 |
| GO:0043062 | extracellular structure organization | 1.45E-03 |
| GO:0030198 | extracellular matrix organization | 1.69E-03 |
| GO:0045229 | external encapsulating structure organization | 2.12E-03 |
| GO:1901607 | alpha-amino acid biosynthetic process | 3.86E-03 |
| GO:0001501 | skeletal system development | 4.04E-03 |
| GO:0061448 | connective tissue development | 5.13E-03 |
| GO:0032963 | collagen metabolic process | 5.16E-03 |

Molecular function
| Go terms | Description | FDR |
| --- | --- | --- |
| GO:0030020 | extracellular matrix structural constituent conferring tensile strength | 9.84E-08 |
| GO:0048407 | platelet-derived growth factor binding | 5.30E-07 |
| GO:0005201 | extracellular matrix structural constituent | 1.01E-05 |
| GO:0003674 | molecular_function | 1.52E-05 |
| GO:0005488 | binding | 1.63E-04 |
| GO:0019838 | growth factor binding | 7.99E-03 |
| GO:0004748 | ribonucleoside-diphosphate reductase activity, thioredoxin disulfide as acceptor | 2.03E-02 |
| GO:0061731 | ribonucleoside-diphosphate reductase activity | 2.29E-02 |
| GO:0005518 | collagen binding | 4.64E-02 |
| GO:0003824 | catalytic activity | 4.67E-02 |

Cellular component
| Go terms | Description | FDR |
| --- | --- | --- |
| GO:0098644 | complex of collagen trimers | 1.88E-12 |
| GO:0005581 | collagen trimer | 1.40E-10 |
| GO:0140151 | solid phase of interstitial matrix | 1.37E-08 |
| GO:0140152 | collagenous component of interstitial matrix | 1.82E-08 |
| GO:0098643 | fibrillar collagen | 1.20E-07 |
| GO:0005583 | fibrillar collagen trimer | 1.44E-07 |
| GO:0005614 | interstitial matrix | 4.47E-07 |
| GO:0031012 | extracellular matrix | 1.53E-05 |
| GO:0030312 | external encapsulating structure | 2.10E-05 |
| GO:0098647 | collagen beaded filament | 2.21E-04 |

## Discussion

In this study, we investigated the effects of the histone deacetylase inhibitor Vorinostat (SAHA) on myogenic differentiation using a multi-omics approach in two different cellular models, C2C12 and L6 cell models. Our findings reveal that HDAC inhibition induces a complex and stage-dependent reprogramming of skeletal muscle cells, affecting proliferation, cytoskeletal organization, metabolism, and extracellular matrix remodeling.

At the functional level, SAHA exerted a modest but consistent reduction in cell viability, with a more pronounced effect in C2C12 compared to L6 cells. Morphological and immunofluorescence analyses further demonstrated that SAHA delays early myogenic differentiation, as evidenced by reduced myotube formation and impaired alignment, while still allowing the formation of multinucleated myotubes at later stages. These observations suggest that HDAC activity is required for the proper timing of myogenic progression rather than for the absolute initiation of differentiation, in line with previous studies investigating other HDAC inhibitors, including trichostatin A (TSA), valproic acid, and butyrate (17,18,21,29,30). Treatment with valproic acid has been reported to enhance myogenic commitment when applied at early stages, particularly by promoting histone acetylation and facilitating the activation of myogenic regulatory factors such as MyoD and myogenin (18). Similarly, HDAC inhibition has been shown to induce differentiation programs and reduce proliferation across multiple cell types, highlighting its role in shifting the balance from proliferation toward lineage commitment (31). Conversely, other studies have demonstrated that continuous or late-stage exposure to HDAC inhibitors, including TSA and valproic acid, can impair differentiation processes, indicating that HDAC activity may act as a critical checkpoint preventing premature or dysregulated differentiation (17,21). In line with this, butyrate, a short-chain fatty acid with HDAC inhibitory activity, has been shown to induce growth arrest and differentiation-associated phenotypes, further supporting the dual role of HDAC inhibition in controlling proliferation and differentiation balance (17).

Our study indicates that, at the transcriptomic level, SAHA induced extensive gene expression changes in both models, with a progressive increase in differentially expressed genes over time. In C2C12 cells, SAHA treatment seems to affect cytoskeletal and muscle structural pathways at early stages, while impacting developmental programs, extracellular matrix remodeling, and translational processes at a later stage.

In L6 cells, SAHA treatment affected pathways associated with cell cycle progression, mitotic processes, and cellular motility at an early stage of differentiation, while altering chromatin organization, cytoskeletal regulation, and metabolic processes at day 7.

Proteomic analyses further revealed both conserved and model-specific effects. In C2C12 cells, SAHA promoted a metabolic shift toward mitochondrial activity and oxidative phosphorylation while downregulating cytoskeletal and contractile components, suggesting a disconnection between metabolic activation and structural maturation. In contrast, L6 cells showed upregulation of muscle contraction-related proteins and downregulation of extracellular matrix components, indicating a more advanced structural maturation and reduced connective tissue associated features. The divergent proteomic responses to SAHA exposure in L6 versus C2C12 cells likely reflect a combination of species-specific differences, distinct differentiation capacities, different baseline proteomic and HDAC expression profiles, differential drug sensitivity, and cell line-dependent epigenetic and signaling contexts that shape downstream transcriptional and protein expression programs.

Importantly, protein-level analyses confirmed that SAHA impairs the expression of key myogenic markers, including Myosin heavy chain, Mybpc1, and Tnnt1, particularly in L6 cells, reinforcing the notion of delayed maturation. At the same time, increased histone acetylation levels in both models confirmed effective HDAC inhibition and directly linked the observed molecular and phenotypic changes to epigenetic modulation (16). Similar observations have been reported for other HDAC inhibitors, such as valproic acid, which has been shown to enhance myogenic specification through increased acetylation levels (18). However, it has been suggested that the effects of HDAC inhibition may not be limited to histone modifications alone, as acetylation of non-histone proteins can also contribute to the regulation of myogenic programs. This broader spectrum of targets may explain the complex and sometimes divergent effects of HDAC inhibitors on muscle differentiation (16). In this context, the transcriptional and proteomic changes observed in our study likely reflect the combined impact of chromatin remodeling and modulation of cytoplasmic and structural proteins, ultimately shaping the differentiation trajectory in a stage-dependent manner. Moreover, the differential response of C2C12 and L6 cells underscores the importance of considering species- and model-specific features when interpreting the impact of HDAC inhibition on skeletal muscle biology.

## Conclusions

Taken together, our findings align with and extend previous observations on HDAC inhibitors, highlighting that their effects on myogenesis are highly dependent on timing, cellular context, and differentiation stage. Early HDAC inhibition may facilitate lineage commitment, whereas sustained exposure appears to disrupt cytoskeletal organization and maturation programs. The differences observed between C2C12 and L6 cells further emphasize the importance of species- and model-specific responses in epigenetic regulation.

Overall, HDAC inhibitors have gained increasing attention for their potential therapeutic application in muscle disorders (32). Notably, Givinostat, a pan-HDAC inhibitor, has been approved by FDA (2024) and EMA (2025) for the treatment of Duchenne muscular dystrophy, where the drug promotes muscle regeneration and reduces fibrosis (11,33,34). This highlights the therapeutic potential of targeting epigenetic regulators in skeletal muscle. Interestingly, our results suggest that the effects of HDAC inhibition are highly dependent on timing, cellular context, and differentiation stage in turn cautioning against directly extrapolating results between different muscle cell models. In this regard, the complex and sometimes contrasting effects observed in C2C12 and L6 cells upon SAHA treatment emphasize the need for precise modulation of HDAC activity to achieve beneficial outcomes without impairing proper myogenic progression. Future studies dissecting the contribution of specific HDAC classes, exposure windows and combination strategies with pro-myogenic cues will be essential to harness the full therapeutic potential of HDAC inhibition in muscle-related diseases.

## Supporting information

C2C12 SAHA

C2C12 control

## List of abbreviations

BP: Biological process
CC: Cellular component
DEGs: differentially expressed genes
DMD: Duchenne muscular dystrophy
ECM: extracellular matrix
GO: Gene Ontology
HATs: histone acetyltransferases
HDACs: histone deacetylases
MF: Molecular function
Mybpc1: Myosin binding protein C1
MyHC: Myosin heavy chain
Tnnt1: Troponin T1
TSA: trichostatin A

## Declarations

### Ethics approval and consent to participate

Not applicable

### Consent for publication

Not applicable

### Availability of data and materials

RNAseq BAM files from control samples have been uploaded in NCBI SRA (accession ID: PRJNA1336407, PRJNA1423193), and BAM files from SAHA-treated samples were deposited under accession ID PRJNA1463610.

## Acknowledgements

We thank Swethaa Natraj Gayathri for the assistance with RNAseq analysis.

## Author contributions

V.S. wrote the original draft. V.S., A.N., M.S. conceived the presented idea. V.S., A.R., A.N., M.S. curated the data. V.S. and A.H. performed the data analysis. J.S., PH.J, B.U., M.S. contributed to the interpretation of these results. SV and AM provided the resource. J.S., PH.J, L.A., B.U., A.N., M.S. supervised the work. A.R., A.H., S.V., A.M., B.U., A.N., M.S. provided the financial support. All authors reviewed the manuscript.

## Supplementary material

Video 1: Video representing twitching of C2C12 control cells at day 6 of differentiation.

Video 2: Video representing twitching of SAHA-treated C2C12 cells at day 6 of differentiation.

