## Supplementary material for "Vorinostat (SAHA) alters the skeletal muscle differentiation program": Supplemnetary Figures

**Supplementary Figures**

**Supplementary Figure S1**

MTT assay

Cell viability (% of control)

|  | CTR C2C12 | SAHA C2C12 | CTR L6 | SAHA L6 |
| --- | --- | --- | --- | --- |
| 24 h | 100 | 88.9658 | 100 | 95.2018 |
| 48 h | 100 | 70.6049 | 100 | 83.427 |
| 72 h | 100 | 68.3074 | 100 | 86.7014 |

Supplementary Figure S1. MTT assay in C2C12 and L6 cells in control and SAHA-treated cells after 24 and 48 h of treatment at the concentration 1 μM.

**Supplementary Figure S2**

**a)**

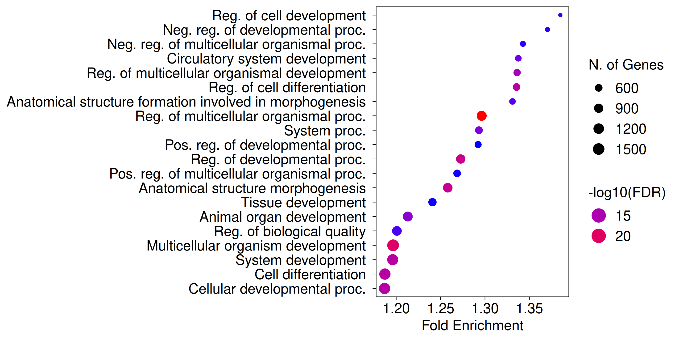

**b)**

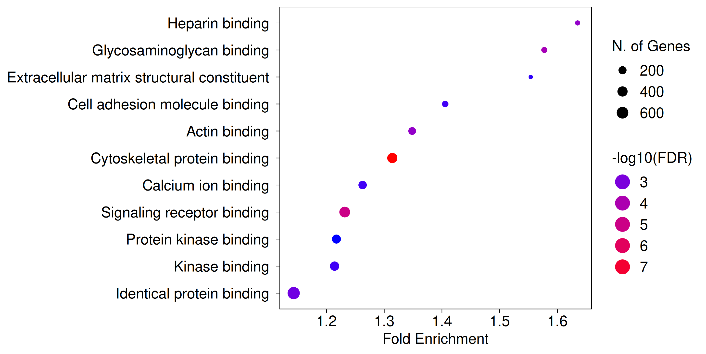

**c)**

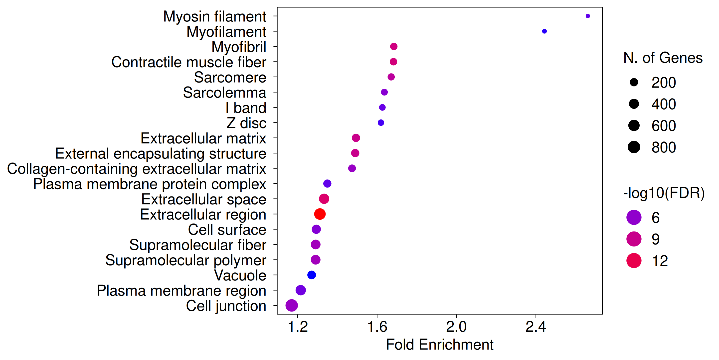

Supplementary Figure S2. ShinyGO Gene Ontology (GO) analysis of C2C12 RNAseq data at day 3 of differentiation. a) Biological process, b) Molecular function, c) Cellular component databases were used to analyse the most enriched pathways associated with DEGs.

**Supplementary Figure S3**

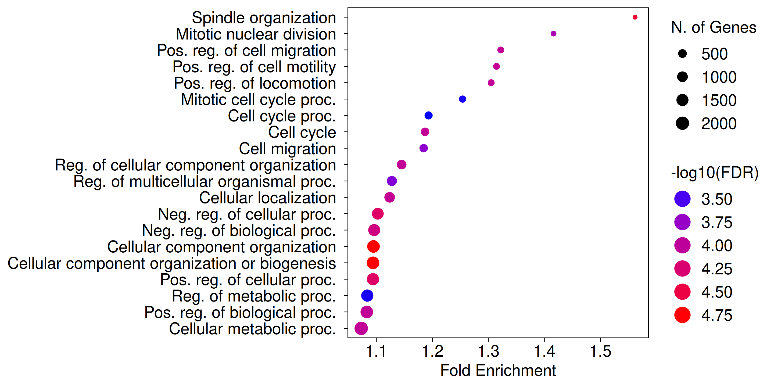

**b)**

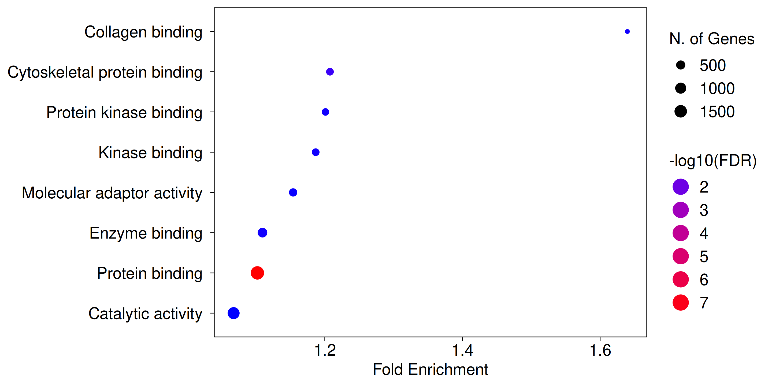

**c)**

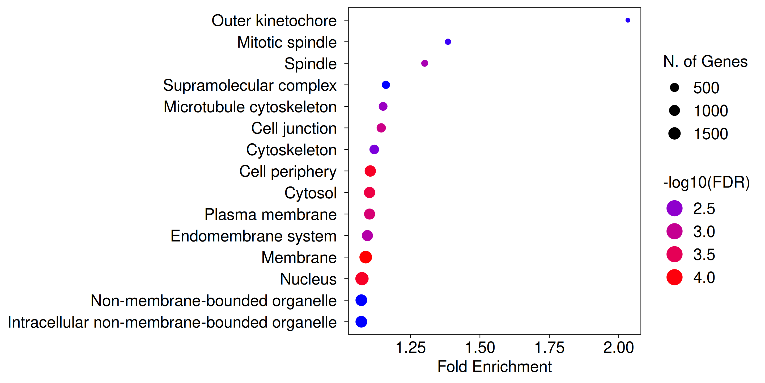

Supplementary Figure S3. ShinyGO Gene Ontology (GO) analysis of L6 RNAseq data at day 3 of differentiation. a) Biological process, b) Molecular function, c) Cellular component were databases were used to analyse the most enriched pathways associated with DEGs.
