## Supplementary Tables for "Vorinostat (SAHA) alters the skeletal muscle differentiation program"

**Supplementary Table**

**Supplementary Table S1.** Group of C2C12biological replicates for RNAseq analysis.

| **Group C2C12** | **Samples** |
| --- | --- |
| CTRDay0 | M1,M2,M4,M5,M6 |
| CTRDay3 | ,M7,M8,M9,M10 |
| CTRDay7 | M19,M20,M21,M22 |
| CTRDay16 | M31,M32,M33,M34 |
| SAHADay3 | M17,M16 |
| SAHADay7 | M27,M28,M29,M30 |
| SAHADay16 | M40,M41,M42 |

**Supplementary Table S2.** Group of L6 biological replicates for RNAseq analysis.

| **Group L6** | **Samples** |
| --- | --- |
| CTRDay0 | M1,M2,M4 |
| CTRDay3 | M5,M6,M7,M8 |
| CTRDay7 | M9,M10,M11,M12 |
| SAHADay3 | M21,M22,M23,M24 |
| SAHADay7 | M25,M26,M27,M28 |
